# A functional different immune capacity in cattle is associated with higher mastitis incidence

**DOI:** 10.1101/311316

**Authors:** Karina Lutterberg, Kristina J. H. Kleinwort, Bernhard F. Hobmaier, Stefanie M. Hauck, Stefan Nüske, Armin M. Scholz, Cornelia A. Deeg

## Abstract

Bovine neonatal pancytopenia (BNP) was a deadly disease transferred by antibodies from 5-10% of cows given a novel BVD vaccine. Disease was lethal in 90% of calves receiving colostrum with BNP antibodies. The cause of BNP is not fully understood to date. We revealed a profound difference in immune capacities between BNP dams and non-responders. Significant differences were detectable in response to *in vitro* stimulation of peripheral blood derived lymphocytes to several mitogens and IL-2. BNP cows regulated their immune proteomes completely different from controls with other immune response master regulators. Since we detected this response pattern also in 22% of cows that were never vaccinated at all, this immune deviant (ID) phenotype is still present in cattle and probably inherited. Immune response pattern of these cows was stable over an observation period of 38 months. Importantly, ID have a significant increased prevalence of mastitis underscoring the clinical importance.

## INTRODUCTION

Vaccines are the most effective and also affordable disease-prevention tools (1) and maternal antibodies protect the offspring from infections in man (2) and animals (3). Pre-partum vaccination of cows with different virulence factors of bacterial and viral infectious diseases effectively stimulates the production of specific antibodies in the colostrum (4, 5). However, maternal vaccination of cows with a new BVD vaccine, which was manufactured by a novel production technology and addition of a highly potent adjuvant, triggered unforeseeable fatal immune reactions with production of pathogenic antibodies in some cows that were then transferred to calves (6, 7).

BNP was a disease of newborn calves with an extremely high lethality rate of 90% (8). Since 2006, BNP first occurred in southern Germany (9) and then in several other countries (10). Affected calves suffered from hemorrhagic diathesis, thrombocytopenia, leukocytopenia and bone marrow depletion, which resulted in a deadly bleeding disorder (10-12). The vaccination with PregSure BVD was proven to be the cause for BNP (6, 13-15). From all vaccinated cows, only 5-10% produced pathogenic BNP alloantibodies (7, 16). The disease was transmitted via colostrum antibodies by this subgroup of Pregsure BVD vaccinated cows to their calves and other, non-related calves that also received colostrum of these BNP dams (7, 10). Other authors postulated that Pregsure BVD vaccination induced alloantibodies in BNP dams and the major histocompatibility complex class I (MHC I) was the dominant alloantigen target (17-20). This hypothesis could not be confirmed (21) and recently, other target candidates were suggested (15, 20). In contrast, there were also open questions about a possible general difference in immune responses between Pregsure BVD vaccinated control cows and BNP donors.

From own analyses, we concluded that the BNP donors could per se substantially differ in their immune capacity, which would then have resulted in this fundamentally different immune response induced through vaccination in these cows. Therefore, we characterized the immune response of BNP donors and controls in this study in-depth and found a markedly different immune capacity in BNP dams. This different immune capacity was readily detectable in 22% of cows in an PregSure BVD unvaccinated cohort, confirming its current existence in cattle. Finally, the functional correlation of the different immune phenotypes with the frequency of several diseases was analyzed.

## RESULTS

### Lymphocytes of BNP dams show hyperproliferation after polyclonal stimulation *in vitro*

After *in vitro* stimulation (48h) with T and B cell mitogen PWM (22) and T cell mitogen ConA (23), a clearly divergent reaction of lymphocytes from PregSure BVD vaccinated cows became evident. Cells from cows known to produce BNP antibodies after vaccination (BNP lymphocytes) proliferated significantly stronger (4.5 fold) than cells from vaccinated control dams after PWM stimulation (Fig. 1A, BNP to Ctr, **** p < 0.0001) and ConA stimulation (8 fold stronger; Fig. 1B, BNP to Ctr, **** p < 0.0001). Thus, *in vitro* proliferation assays revealed a highly significant hyperproliferation of BNP lymphocytes demonstrating an increased reaction to polyclonal immune stimulation in these cows.

**Figure 1:**
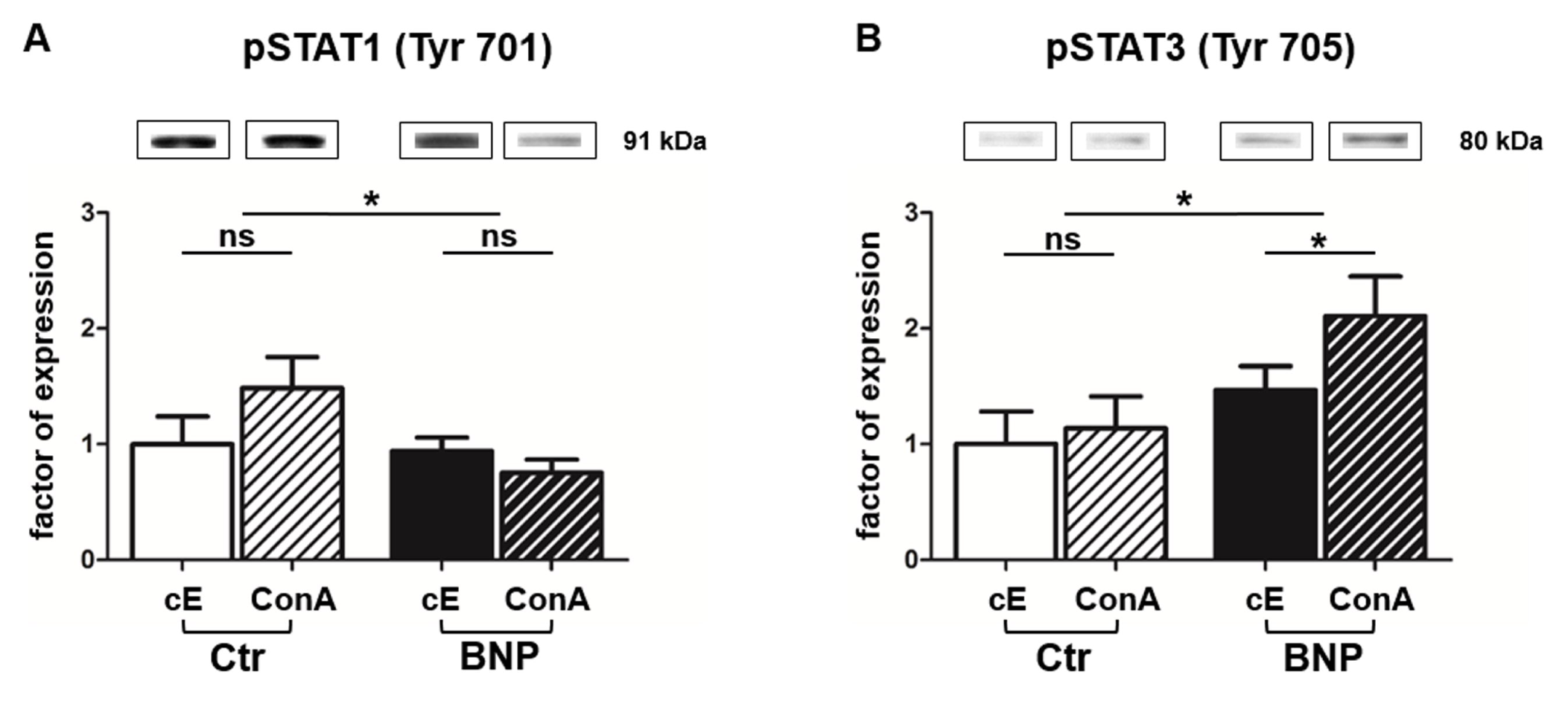
Hyperproliferation of BNP lymphocytes after polyclonal stimulation *in vitro*. (A) Lymphocytes from BNP cows (black bars, n = 5, technical replicates n = 98) proliferated 4.5 times stronger (**** p < 0.0001 vs. control) after PWM stimulation than lymphocytes from control dams (white bars, n = 6, technical replicates n = 53). (B) Increased proliferation rate (8 fold; **** p < 0.0001 vs. control) of lymphocytes from BNP dams (black bars, n = 5, technical replicates n = 86) compared to control dams (white bars, n = 6, technical replicates n = 52) after *in vitro* stimulation with ConA. Proliferation rate shown in mean ± SD. Mann-Whitney test was used.

### Lymphocyte proteomes reveal differential protein regulation between normal and pathological immune responders to PregSure BVD vaccination

To clarify whether the observed hyperproliferation was just caused by a quantitative difference of the same immune response or by a functional difference, we executed a differential proteomics experiment to discover the immune proteomes and their regulation. We detected 3689 differently regulated proteins after PWM stimulation of lymphocytes and 5471 after ConA stimulation. Hierarchical cluster analysis of respective proteomes already revealed fundamental differences in protein abundances between control and BNP cows in constitutive protein levels (cE, Fig. 2 A/B). Differences in protein regulation were even stronger after immune stimulation (PWM and ConA, Fig. 2 A/B, hierarchical clustering of all identified proteins). These proteome analyses therefore revealed substantial quantitative and qualitative differences on protein level.

**Figure 2:**
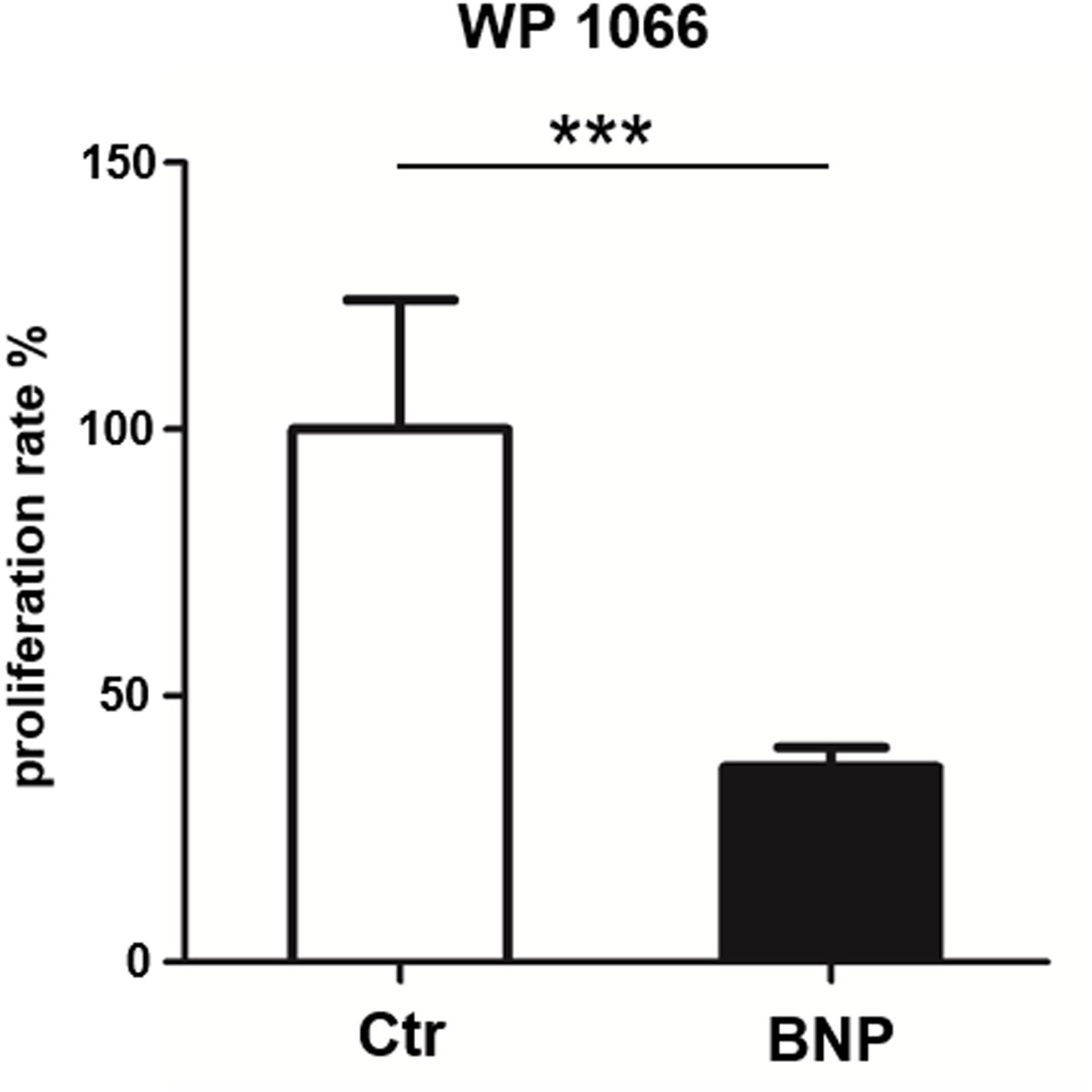
Lymphocyte proteomes reveal regulation of differential proteins between normal and pathological immune responders to PregSure BVD vaccination. Differential proteome analyses of PBL from control (Ctr, n = 2) and BNP (n = 2) cows already revealed differences in protein expression patterns in constitutive expression (cE) of proteins, that shifted even stronger after (A) PWM and (B) ConA stimulation (48h). Highly abundant proteins are presented in green and low abundant proteins in red. (C/D) List of all identified STAT proteins in bovine PBL after polyclonal activation of lymphocytes with PWM (C) or ConA (D). STAT3 and STAT5A were selectively regulated in BNP lymphocytes (C) compared to controls due to PWM. After ConA stimulation, controls significantly upregulated STAT1 and STAT6 (D), whereas in PBL of BNP cases, STAT3 was enhanced during immune response (D).

### Immune stimulation leads to different usage of STAT pathways in PBL of control and BNP cows

Next, we analyzed transcription pathways in PBL after polyclonal stimulation in control and BNP cows to identify possible differences in master transcription factor expression important for different lymphocyte pathways. Signal transducer and activator of transcription (STAT) induces different T helper (Th) answers in mice and man (24) and therefore we characterized usage of different STAT pathways as a response to immune stimulation in both cow groups. After polyclonal activation of lymphocytes with PWM, STAT3 and STAT5A were selectively upregulated in BNP lymphocytes (Fig. 2C). At the same time, we could not detect higher abundance of any STAT analyzed in controls. In contrast, as reaction to ConA stimulation, controls significantly upregulated STAT1 (Fig. 2D), whereas in PBL of BNP cases, STAT3 was enhanced in immune response (Fig. 2D).

### Lymphocytes of control and BNP cows regulate different STATs in response to ConA stimulation *in vitro*

We decided to further analyze the differential activation of STATs to the T cell mitogen ConA, we determined phosphorylation of STAT1 and STAT3 in response to immune stimulation. In lymphocytes of controls, phosphorylation of STAT1 Tyr701 significantly increased after *in vitro* stimulation with ConA (Fig. 3A, Ctr to BNP after ConA stimulation, * p < 0.05). In contrast, lymphocytes of BNP dams regulated the immune response through phosphorylation of STAT3 Tyr705 (Fig. 3B, BNP to Ctr, * p < 0.05). These experiments ascertained the qualitative difference in immune reactions between the cow groups to a T cell stimulus.

**Figure 3:**
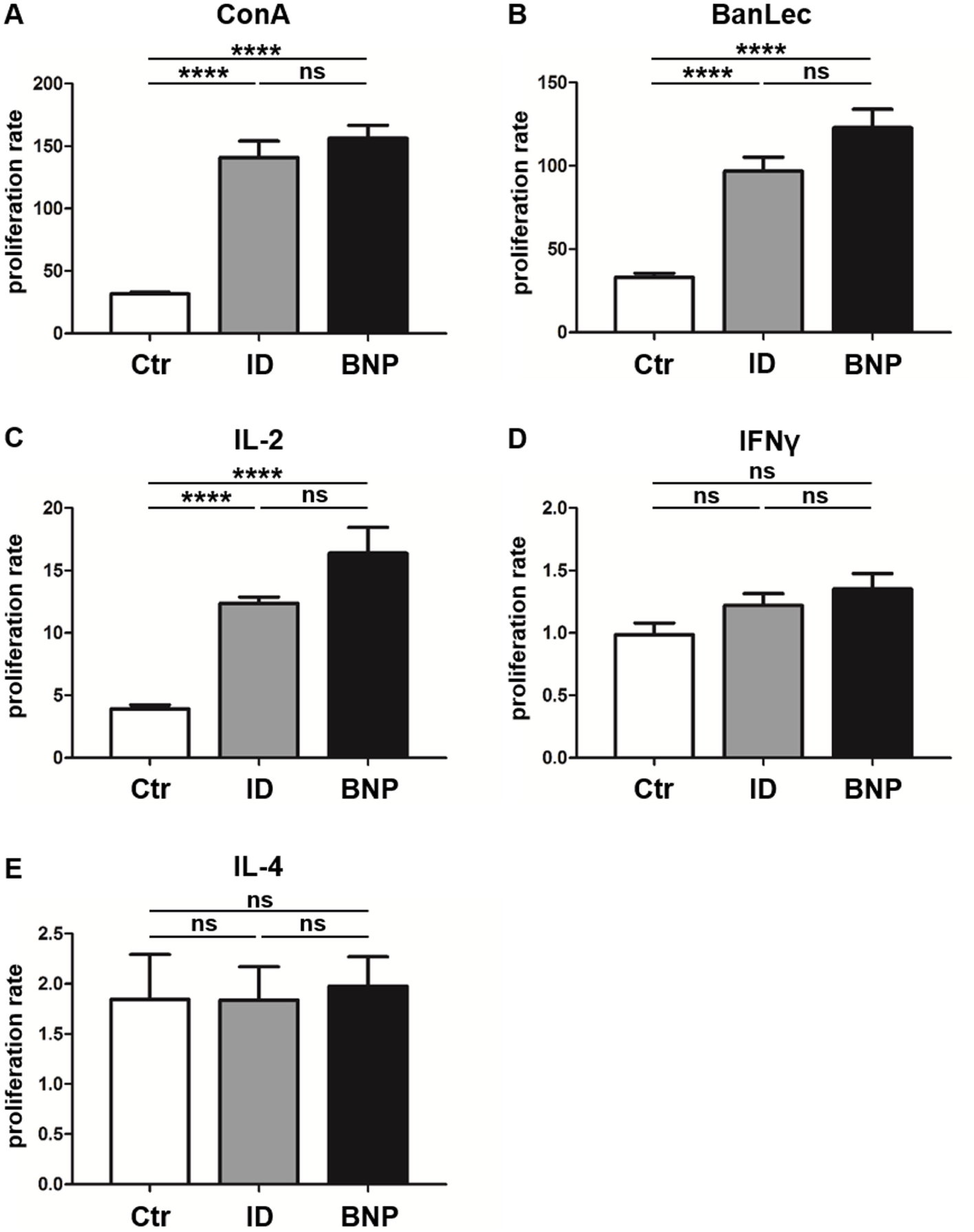
In PBL of control and BNP cows, different transcription factors were activated after ConA stimulation *in vitro*. (A) Control lymphocytes (white-black striped bars, n = 5) phosphorylated STAT1 significantly stronger than BNP lymphocytes (black-white striped bars, n = 5; * p < 0.05) after ConA stimulation. (B) Only BNP lymphocytes activated STAT3 in response to ConA stimulation (* p < 0.05). Protein expression is shown in mean ± SD. For statistically analyses Student’s *t* test was performed.

### Inhibition of STAT3 in PBL of BNP cows abolished hyperproliferation to polyclonal immune stimulation

Since BNP lymphocytes responded to polyclonal stimulation with hyperproliferation and activation of STAT3, we next tested if inhibition of STAT3 reversed respective hyperproliferation. WP1066 (STAT3 inhibitor) significantly reduced proliferation of BNP lymphocytes after ConA stimulation (Fig. 4, inhibition of BNP to Ctr, *** p < 0.001), but not of control lymphocytes. This verified the importance of STAT3-pathway for the hyperproliferation in BNP PBL after polyclonal T cell stimulation.

**Figure 4:**
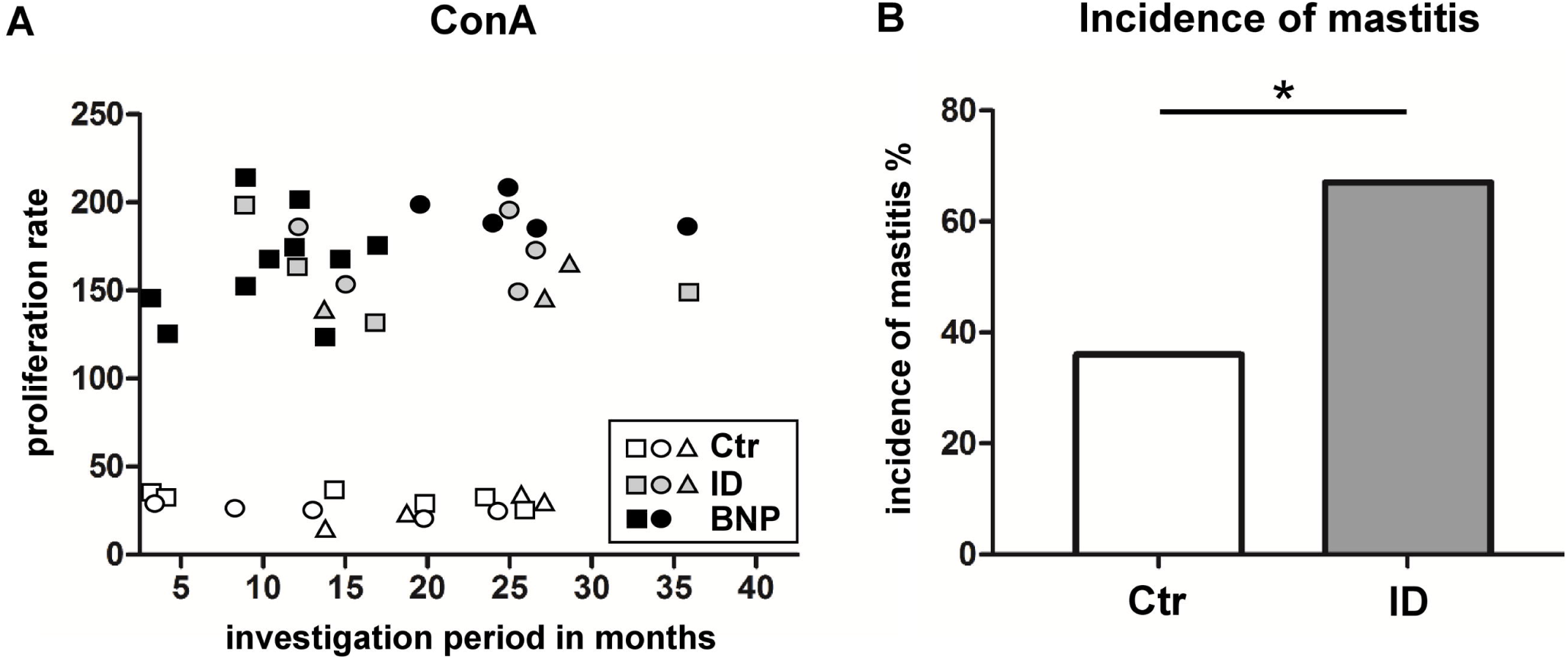
STAT3 inhibitor WD1066 selectively inhibits hyperproliferation of lymphocytes from BNP cows. STAT3 inhibitor did not inhibit proliferation of control PBL (white bar, n = 4), but significantly inhibited ConA stimulated BNP cells (black bars, n = 4, *** p < 0.001). Percentage of inhibition rate shown in mean ± SD. Student’s *t* test was used for statistical analyses.

### Immune deviant phenotype is also detectable in unvaccinated cows

Since we first detected the deviant immune phenotype in BNP cows, we were next interested to see if we could find evidence for this immune phenotype in cows that were never vaccinated with PregSure BVD. This was important to clarify whether respective phenotype was induced through the vaccination or if it already existed in some cows before. The latter would mean that this immune capacity is probably congenital. In order to test this, we examined response of lymphocytes to different polyclonal stimuli in a large group of unvaccinated cows from one farm, where a difference due to environmental factors could be excluded. In *in vitro* proliferations assays with T cell mitogen ConA we observed that 22% of the unvaccinated cows reacted similar to BNP dams. The tests showed a hyperproliferative reaction to ConA (immune deviant (ID) phenotype; Fig. 5A, reaction difference ID to controls or BNP to controls, **** p < 0.0001). Lymphocytes from ID cows with a hyperproliferative immune phenotype reacted with no significant difference to lymphocytes of BNP cows in these assays (Fig. 5A, reaction difference BNP to ID).

**Figure 5:**
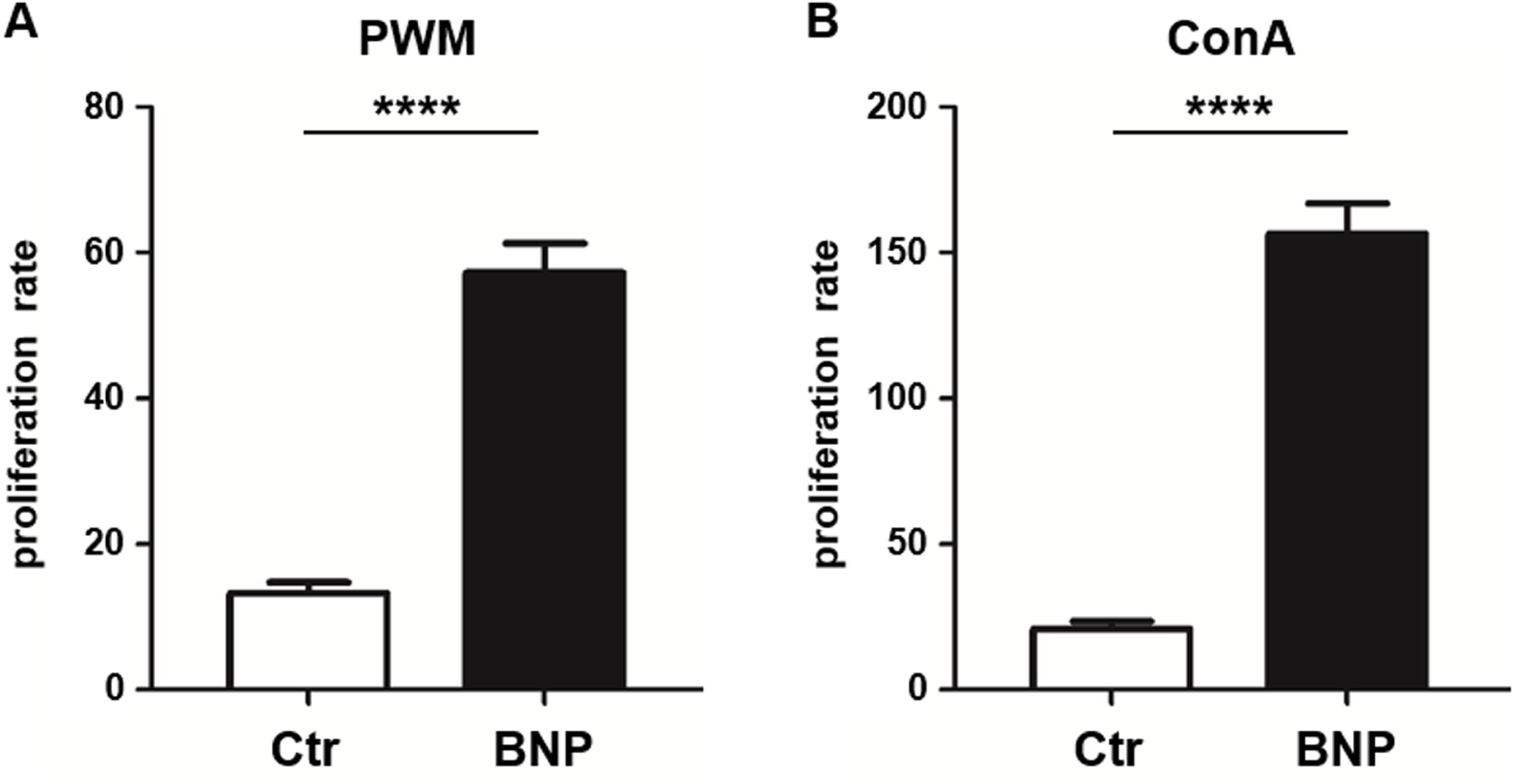
Immune deviant phenotype also detectable in unvaccinated cows. (A) In *in vitro* proliferation assays, 22% of unvaccinated cows showed a BNP-like hyperproliferative reaction to ConA (Fig. 5A, **** p < 0.0001 vs. Ctr (white bar, n = 121)). Lymphocytes from ID cows (grey bars, n = 27) with a hyperproliferative immune phenotype showed no significant (ns) differences to lymphocytes of BNP cows (black bars, n = 4, technical replicates n =86). (B) After BanLec stimulation, ID lymphocytes reacted hyperproliferative in comparison to control lymphocytes (**** p < 0.0001) and the proliferation rates of these ID lymphocytes did not show significant differences (ns) compared to BNP lymphocytes. (C) After IL-2 stimulation, lymphocytes of ID cows also reacted excessively (**** p < 0.0001 vs. Ctr) just like lymphocytes from BNP donors (ns). (D/E) After IFNγ (D) and IL-4 (E) stimulation, no significant differences between lymphocytes of unvaccinated and BNP cows were determined (ns). Proliferation rate shown in mean ± SD. Student’s *t* test was used.

### Lymphocytes of ID cows also react to BanLec and IL-2 with hyperproliferation

In order to further characterize the pathway targeted by the polyclonal activators PWM and ConA in the cows we tested additional mitogens. The T cell mitogen BanLec (24), which activates T cells of man through the IL-2 pathway (25, 26), led to similar hyperproliferation of PBL from BNP dams as observed with ConA (Fig. 5A, reaction difference factor: 5.0, BNP to controls, **** p < 0.0001). PBL of ID cows responded to BanLec stimulation with a similar immune response intensity as BNP (Fig. 5B, reaction difference factor: 3.0 ID to controls, **** p < 0.0001). Additionally, we detected no significant difference in the reaction of ID and BNP lymphocytes. Since the results with ConA and BanLec pointed to a response via IL-2 pathway, we next tested stimulation with purified bovine IL-2 only. After IL-2 stimulation, lymphocytes of ID cows also reacted excessively (Fig. 5C, reaction difference factor: 3.1, ID to controls, **** p < 0.0001) and did not show significant differences to BNP lymphocytes (Fig. 5C, reaction difference factor: 3.7, BNP to controls, **** p < 0.0001).

### IL-2, but not IFNγ or IL-4 promote the different immune response in ID cows

Finally, we tested further signature cytokines for different Th immune responses, Interferon gamma (IFNγ; Th1) and Interleukin 4 (IL-4; Th2), to analyze differentiation of T helper subsets. We detected no significant differences between lymphocytes of vaccinated and unvaccinated controls and BNP cases after IFNγ or IL-4 stimulation *in vitro* (Fig. 5D/E). This proves a crucial role for IL-2 in the deviant immune responses, but not for IFNγ or IL-4. We discovered a significant proportion of cows that responded like BNP-dams, but have never received PregSure BVD vaccine. The cytokine IL-2 plays an essential role for the dysregulation in bovine lymphocytes of these immune deviant animals.

### Immune deviant phenotype in cows correlates with significantly increased incidence of mastitis

In this study, the proliferative phenotype of controls and ID cows could be repeatedly tested during an overall observation period of 38 months. Interestingly, retested animals reacted consistently in the *in vitro* proliferation assays (Fig. 6A, proliferation rate after ConA stimulation, data of three representative controls, ID cases and two BNP cows, sampled at least three times). PBL of controls (Fig. 6A white symbols) reacted stable in the assay, as well as the PBL of hyperproliferative animals (ID and BNP), that were steadily hyperproliferative (Fig. 6A grey and black symbols).

**Figure 6:**
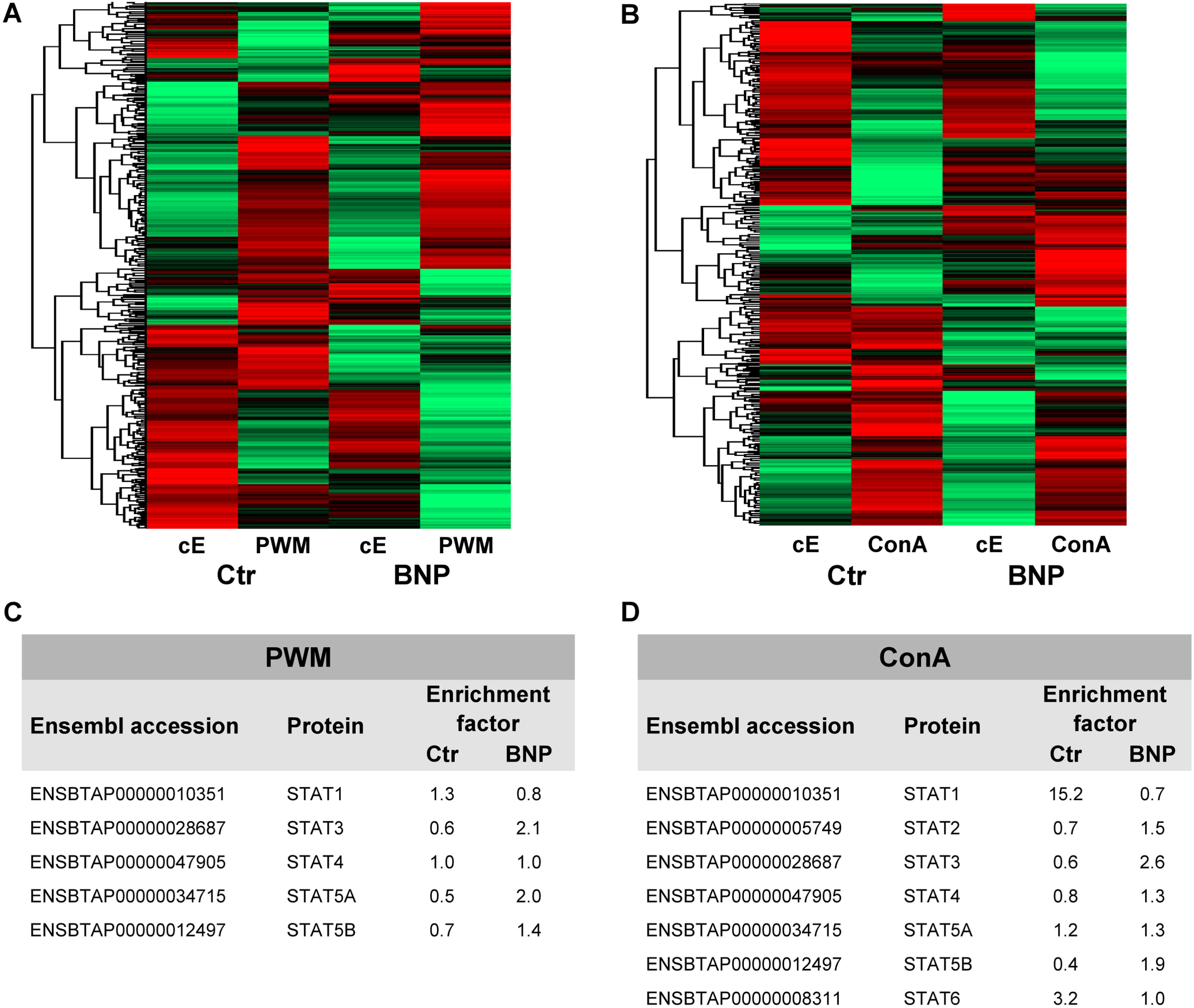
Proliferation response types remain consistent and ID have a significantly increased incidence of mastitis. (A) The hyperproliferative phenotype of BNP cows (black) and ID cows (grey) as well as the low proliferation of control cows (white) were consistently detectable over the entire observation period (38 months). Three representative animals per group are shown, white symbols=controls, grey symbols=ID, black symbols=BNP. (B) ID cows (grey, n = 27) suffered twice as often (difference in incidence * p < 0.05) from mastitis than the control cows (white, n = 73). Incidence of mastitis shown in percent. Odds ratio and chi-square distribution was used for statistical analyses.

Since we wanted to know if this newly described immune deviant phenotype related to impaired health parameters in respective cows, we analyzed milk production and health parameters of control and ID cows of the same farm. From a total of 16 parameters tested, there was no difference in 15 parameters. But, mastitis incidence was significant different between both cow groups. We detected that ID cows suffered twice as often as control cows from mastitis (Fig. 6B, * p < 0.05). This clearly indicates that the hyperproliferative phenotype of ID cows has further and important clinical relevance apart from the unwanted response to the BVD vaccine.

## DISCUSSION

BNP demonstrated that an unwanted, fatal immune response occurred in a group of 5-10% cows of the animals vaccinated with PregSure BVD (7, 16). These BNP dams developed aggressive antibodies that killed 90% of calves that received these antibodies via colostrum, irrespective if they were related to the dam or if they received the antibodies via mixed colostrum (8, 10). It is still unclear, why a subgroup of cows responded differentially to the vaccination. We hypothesized that the vaccine met some cows that per se differed significantly in their immune capacity. This conjunction led in turn to the production of pathologic antibodies in these BNP dams.

In this study, we first detected a significantly different proliferative response in PBL of BNP dams (hyperproliferation) after polyclonal *in vitro* stimulation with two different mitogens, confirming a quantitative difference in immune activation of BNP PBL (Fig. 1A/B). Next, we analyzed the constitutive PBL proteomes and their regulation after polyclonal immune stimulation in order to investigate a potential functional difference in the immune response. Proteome analyses revealed a total of 5471 proteins expressed in bovine lymphocytes. The proteomics approach already substantiated a qualitative difference in constitutive expression of proteins in PBL of controls and BNP dams (cE, Fig. 2A/B). After immune stimulation, the regulated proteins (factor ≥ 2.0) differed even further between controls and BNP, confirming significant qualitative differences in immune responses.

Since different STATs initiate the differentiation of various Th cell subsets (24) as first response master regulators, we subsequently analyzed different STATs in the cells after immune stimulation as indicators of immune response pathway choice. STATs provide lineage specificity by promoting the differentiation of a given Th cell subset while opposing the differentiation to alternative Th cell subsets (27).

In control lymphocytes an upregulation (Fig. 2D) and a significantly higher activation (Fig. 3A) of STAT1 after T cell stimulation was found. The effect of STAT1 inhibition could not be tested, since we found no STAT1 specific inhibitor that was commercially available. In mice and man, STAT1 is important for the differentiation of naive CD4^+^ T cells to Th1 cells (28) or to type 1 regulatory (Tr) T cells (29). From these findings, we conclude that after immune stimulation, control lymphocytes use the STAT1 pathway and we hypothesize that control cows react with a Th1 or Tr immune response in polyclonal stimulation assays.

In contrast, STAT3 was upregulated in PBL proteome of BNP dams (Fig. 2D) and also showed increased phosphorylation (Fig. 3B) in BNP lymphocytes after immune stimulation. This is an interesting finding, since high STAT3 expression level is associated with autoimmune disease in man (30). STAT3 could therefore be a possible master regulator for the deviant immune response in BNP dams. In mouse and man, STAT3 induces the development of follicular Th (Tfh), Th17 and Th22 cells (27, 31, 32). Our findings proved a STAT3 pathway dependent immune response of BNP lymphocytes after polyclonal stimulation, but so far it is unknown which Th subsets are regulated by STAT3 in cows. The dependence of the hyperproliferative phenotype of immune deviant PBL on STAT3 was shown by inhibition of respective factor using WP 1066, a cell permeable tyrphostin analog which blocks the STAT3 pathway through inhibition of Janus kinase 2 (JAK2) protein tyrosine kinase (33). As a result, we observed a significantly reduced proliferation to 37% in BNP PBL, but no effect in control PBL (Fig. 4). Our findings therefore show, that the deviant immune response of BNP donors is associated with STAT3/JAK2 pathway. In further studies, we will clarify whether immune cells from control cows differentiate to Th1 or Tr subsets and BNP donor cows develop Tfh, Th17 or Th22 cells after immune stimulation.

In order to confirm the hypothesis of the deviant immune phenotype detected in BNP cows being present in these animals even before PregSure BVD vaccination, the immune capacities of non-PregSure BVD vaccinated cows were analyzed along the same lines as BNP characterization. To exclude further influencing factors, 121 cows from one dairy farm were tested and health data were examined in a retrospective study for 38 months. We observed that 22% of unvaccinated cows (ID cows) showed a BNP-like hyperproliferative response to T cell stimulation *in vitro* (Fig. 5A/B). All 27 immune deviant cows, sampled at least three times, showed consistent hyperproliferative reaction to ConA (Fig. 6A). The ID animals were always significantly different in their immune response compared to the control cows (Fig. 6A). We could thus prove, that a certain percentage of the cattle population has an immune deviant phenotype per se, and that this phenotype was not caused by vaccination with PregSure BVD.

The immune deviant response described here clearly depends on IL-2. Since BanLec activates T cells in man (24, 26) via the IL-2 pathway (25), the hyperproliferative response of BNP and ID animals to BanLec (Fig. 5B) indicated a role for IL-2. This was confirmed by sole application of IL-2 (Fig. 5C). The specific function of IL-2 for bovine T cells was not described so far. In man and mice, IL-2 has pleiotropic autocrine or paracrine activities that regulate proliferation of T cells (34). But it was also shown to play a key role in promoting development, homeostasis and function of regulatory T cells though IL-2/STAT5 signals (35). Further, IL-2 balances expression of different STATs in Th cells of mice (36). Therefore, the association of the immune deviant response in cows with IL-2 is very interesting and merits further in-depth investigations in future studies.

To determine, whether the newly detected immune phenotype is of clinical relevance, we analyzed 16 clinical parameters in 100 cows of the sampled dairy farm and compared the incidences between controls (n=73) and ID animals (n=27). Interestingly, there was a significant difference in one health parameter: mastitis (Fig. 6B). ID cows had a significantly higher occurrence of mastitis than control cows, providing an important correlation between deviant immune phenotype and mastitis incidence (Fig. 6B). Future studies will focus on the context enhanced mastitis incidence in the newly discovered deviant immune phenotype. The hypothesis is an altered immune reaction of ID cows to classical mastitis pathogens, facilitating mastitis onset in these animals.

In conclusion, we demonstrated hyperproliferation by BNP lymphocytes, which prefer a STAT3/JAK2 regulated pathway after immune stimulation. IL-2 plays an important role in divergent immune response of ID lymphocytes. Further, we detected 22% of non-PregSure BVD vaccinated cattle with hyperproliferative phenotype in the cohort investigated by us. That means that the immune deviant phenotype exists independently of the vaccination in recent cattle. Their deviant immune response pattern is identical to BNP cows. IL-2 plays an important role in divergent immune response of ID and BNP lymphocytes. Importantly, we discovered a correlation of the immune deviant phenotype with high mastitis incidence.

## MATERIALS AND METHODS

In this study, PBL of a total of 132 cows were used. Six were PregSure BVD-vaccinated control cows, which did not produce pathologic antibodies and five were confirmed BNP dams that were vaccinated with PregSure BVD, that had calves which died from hemorrhagic diathesis and had confirmed thrombocytopenia, leukocytopenia and bone marrow depletion (10). Additionally, 121 healthy cows (female, age: two to ten years, first to eight lactation) of one dairy farm were tested.

### Proliferation assays

PBL were seeded in 96-well plates (1 × 10^5^ cells/well) and stimulated (36h or 48h w/o inhibition) with either PWM, ConA or BanLec (Sigma-Aldrich, Taufkirchen, Germany, 5µg/ml), bovine IL-2 (Bio-Techne, Wiesbaden, Germany, 1ng/ml), bovine IL-4 or human IFNγ (Biomol, Hamburg, Germany, 1ng/ml). For the inhibition assays, stimulation was followed by STAT3 inhibitor (WP 1066, Santa Cruz, Heidelberg, Germany, 50ng/ml, 12h). After incubation, the cells were pulsed for 14h with 0,05 mCi/well [methyl-^3^H]-thymidine (Perkin Elmer, Hamburg, Germany), harvested and counts per minute were measured. Proliferation rate after stimulation was expressed as factor of [^3^H]-thymidine incorporation with respect to the unstimulated cells or after inhibition as percentage to ConA stimulated cells.

### Differential proteome analyses

PBL (2.2 × 10^7^ cells) of two PregSure BVD vaccinated control cows and two BNP dams were stimulated with PWM and ConA (5µg/ml) for 48h. For LC-MS/MS analysis, peptides were separated on a reversed chromatography column (75µm IDx15cm, Acclaim PepMAP 100 C18. 100Å/size, LC Packings, Thermo Fisher Scientific, Bremen, Germany) and the analysis was conducted with an Ultimate 3000 nano-HPLC system (Dionex, Idstein, Germany). Nano-HPLC was connected in a linear quadruple ion trap-Orbitrap (LTQ Orbitrap XL) mass spectrometer (Thermo Fisher Scientific, Bremen, Germany). The mass spectrometer was operated in the data-dependent mode to automatically switch between Orbitrap-MS and LTQ-MS/MS acquisition. Up to the ten most intense ions were selected for fragmentation on the linear ion trap using collision induced dissociation at a target value of 100 ions and subsequently dynamically excluded for 30s. Elevated spectra were imported into Progenesis software (version 2.5). After comparison and normalization, spectra were exported as Mascot Generic files and searched against the Ensembl bovine database (version 80) with Mascot (Matrix Science, version 2.4.1). Peptide assignment was reimported to Progenesis Software. All unique peptides allocated to a protein were considered for quantification. Only proteins quantified with at least two peptides were included for further analysis. For hierarchical cluster analyses, all identified proteins were described in the Perseus software (version 1.6.1.1).

### Western Blots

For protein expression analyses with Western blots, PBL were first lysed in lysis buffer (9 M Urea, 2 M Thiourea, 65 mM Dithioerythritol, 4% CHAPS). Proteins were then separated by SDS-PAGE on 8% gels (7 μg protein/slot) and blotted semidry onto 8.5 × 6 cm PVDF membranes (GE Healthcare, Freiburg, Germany) and blocked with 4% BSA (1h). Blots were incubated overnight with respective primary antibodies: rabbit anti-pSTAT1 Tyr701, rabbit anti-pSTAT3 Tyr705 (Cell Signaling, Darmstadt, Germany, 1:500) or mouse anti-beta actin (Sigma, Taufkirchen, Germany, 1:5000). As secondary antibodies, either HRP-coupled goat anti-rabbit IgG (H+L) antibody (Cell Signaling, Darmstadt, Germany, 1:5000) or goat anti-mouse IgG (H+L) antibody (Sigma-Aldrich, Taufkirchen, Germany, 1:5000) were used (1h). Signals were detected by enhanced chemiluminescence on X-ray film (SUPER-2000G ortho, Fuji; Christiansen, Planegg, Germany). Films were scanned on a transmission scanner and densitometric quantification of Western blot signals was performed using ImageJ software (open source: http://imagej.nih.gov/ij/). Abundances of pSTAT1 and pSTAT3 were subsequently normalized to beta actin.

### Health parameters

Milk production and health parameters (observed by veterinarians) of 100 non-vaccinated cattle from one dairy farm were analyzed for 38 months retrospectively and correlated to immune response data of *in vitro* assays. 73 control cows with low proliferation rate after ConA stimulation and 27 ID cows, which showed a BNP-like hyperproliferation to ConA were included in this study. Sixteen parameters were analyzed: daily milk yield, average lactation performance (300 days), milk structure (lactose, fat, urea, somatic cell count), fertility parameters (amounts of inseminations, calving-to-conception intervals, medicinal induction of oestrus, ovarian cysts), diseases of musculoskeletal system, claws, digestive tract, respiratory system and metabolic disorders (ketosis, hypocalcemia). All parameters were recorded and listed by the same veterinarians and the statistical analysis was performed using the odds ratio and chi-square distribution.

### Statistical Analyses

Data were analyzed in Prism software (GraphPad, version 5.04) with Kolmogorow-Smirnow (KS) test. If KS test was significant (p < 0.05; normal distribution), Student’s *t*-test was used for statistical analysis, if KS test was not significant (p > 0.05; no normal distribution) statistics were performed using Mann-Whitney test. For statistical analysis of health parameters, we used odds ratio and chi-square distribution. Differences were considered statistically significant with the following p values: * p < 0.05, ** p < 0.01, *** p < 0.001, and **** p < 0.0001.

## Acknowledgements

The authors thank Sven Reese for statistical analyses and Roxane Degroote for critical discussions.

## Author Contributions

C.D. conceived and designed the experiments; K.L., B.H. and K.K. performed the experiments; K.L., K.K., S.H., S.N., A.S. and C.D. analyzed the data; K.L. and C.D. wrote the manuscript. All authors critically read the manuscript and approved the final version to be published. The authors declare no conflict of interest.

